# A human adenovirus encoding IFN-γ can transduce Tasmanian devil facial tumour cells and upregulate MHC-I

**DOI:** 10.1101/2022.05.29.493930

**Authors:** Ahab N. Kayigwe, Jocelyn M. Darby, A. Bruce Lyons, Amanda L. Patchett, Leszek Lisowski, Guei-Sheung Liu, Andrew S. Flies

## Abstract

The devil facial tumour disease (DFTD) has led to a massive decline in the wild Tasmanian devil (*Sarcophilus harrisii*) population. The disease is caused by two independent devil facial tumours (DFT1 and DFT2). These transmissible cancers have a mortality rate of nearly 100%. An adenoviral vector-based vaccine has been proposed as a conservation strategy for the Tasmanian devil. This study aimed to determine if a human adenovirus serotype 5 could express functional transgenes in devil cells. As DFT1 cells do not constitutively express major histocompatibility complex class I (MHC-I), we developed a replication-deficient adenoviral vector that encodes devil interferon gamma (IFN-γ) fused to a fluorescent protein reporter. Our results show that adenoviral-expressed IFN-γ was able to stimulate upregulation of beta-2 microglobulin, a component of MHC-I, on DFT1, DFT2, and devil fibroblast cell lines. This work suggests that human adenoviruses can serve as vaccine platform for devils and potentially other marsupials.

## Introduction

The Tasmanian devil (*Sarcophilus harrisii*; referred to as ‘devil’ hereafter) is listed as an endangered species due to a massive disease-induced decline in its wild population. The devil facial tumour disease (DFTD) is caused by two genetically independent transmissible cancers that are spread through social interactions. Devil facial tumour 1 (DFT1) was first observed in 1996 and DFT2 was detected in 2014 [1, 2]. DFT1 is widespread across the island state of Tasmania, resulting in an 80% decline in the wild devil population [3, 4]. To date, DFT2 has only been detected in southern Tasmania [2, 5]; spread of DFT2 throughout Tasmania would pose another major threat to the wild devil population.

The mortality rate of both DFT1 and DFT2 is almost 100%. A primary immune evasion pathway of DFT1 cells is via downregulation of cell surface major histocompatibility complex class I (MHC-I) molecules [6]. This allows DFT1 cells to effectively hide from cancer-killing CD8+ T cells. Previous research used DFT1 cells treated with IFN-γ to upregulate MHC-I as the basis for an experimental vaccine [7]. Priming healthy devils with a killed DFT1 cell vaccine led to signs of improved immune activation and recognition of DFT1 cells in laboratory and field trials, but did not prevent devils from developing DFTD [7–9]. However, injecting live IFN-γ treated DFT1 cells into the tumours was able to induce tumour regressions in devils that had been primed with the vaccine [7]. Encouragingly, rare cases of natural DFT1 regressions have been documented in the wild [10–13]. These DFT1 regressions demonstrate that with proper stimulation, the devil immune system can recognise and kill DFT1 cells.

Due to the limited ability of the whole cell vaccine strategy to prevent development of DFTD and the logistical problems of trapping and injecting devils in the wild, an adenovirus-based oral bait vaccine (OBV) has been proposed to protect devils from DFTD [14]. Vaccines distributed in edible baits have been employed to control infectious diseases in wildlife in their natural environments. About 665,000,000 oral bait vaccines (OBV) for rabies were distributed in Europe between 1978 and 2014, resulting in successful elimination of rabies in foxes in more than ten European nations (reviewed in [15]). An adenoviral-based rabies vaccine has been established as a safe and effective rabies control strategy in North America [16–18]. In addition to the prophylactic potential of adenoviral vaccines, in humans adenovirus-mediated delivery of IFN-γ has been used as antitumour immunotherapy [19–21].

Human adenoviruses have been detected in marsupials [22], but the ability of human adenoviruses to transduce marsupials cells or more generally infect marsupials is largely unknown [23–26]. We hypothesized that human adenovirus serotype (Ad5) can transduce cells of Tasmanian devil origin and express transgenes, thus serving as a potential DFT1/2 vaccine platform. To test this hypothesis, we generated a replication-deficient human Ad5 vector that encodes devil IFN-γ fused to a blue fluorescent protein (BFP) reporter. Our results show that a replication-deficient Ad5 vector can transduce devil cell lines and expresses functional IFN-γ.

## Materials and methods

### Cell lines for adenoviral infections

The viral vector transduction was tested using DFT1-C5065, DFT1-1426, DFT1-4906, DFT2-JV, DFT2-SN cell lines, and the devil fibroblast cell line FBB-TD602 [6, 27]. Transduction was also evaluated using human lung carcinoma A549 (ATCC, CCL-185), canine fibroblast tumour A-72 (ATCC, CRL-1542) [28], and Chinese hamster ovary CHO-K1 (ATCC, CCL-61) cell lines. Prior to infection with the virus, the cells were maintained in RPMI 1640 medium (Gibco, 11875-093), 10% heat-inactivated foetal bovine serum (FBS) (Bovogen Biologicals, SFBS), 10 mM HEPES (Gibco, 15630-080), 1% non-essential amino acids (Gibco, 11140-050), 1% sodium pyruvate (Gibco, 11360-070), 1% Antibiotic-Antimycotic (Gibco, 15240-062), and 0.1 mM 2-mercaptoethanol (Gibco, 21985-023) in a humidified incubator at 37°C and 5% CO_2_.

### Preparation of replication-deficient adenoviral vector (advAF300)

We developed a replication-deficient adenoviral vector that carries a fusion gene of codon-optimised devil interferon-gamma (IFN-γ; XM_012550654.1) and mTag-blue fluorescent protein (mTagBFP aka BFP; **Figure 1**) [29, 30]. We used a rigid linker between IFN-γ and the BFP reporter as previously described [30, 31]. Codon optimised synthetic DNA for the IFN-γ-linker-BFP fusion gene was ordered from Genscript (Piscataway, New Jersey) and cloned into the NotI and EcoRV restriction sites in the pShuttle-CMV plasmid (Addgene, 16403). The resulting pAF300 plasmid was linearized with *Pme*I (NEB, R0560S) and then transformed into *Escherichia coli* BJ5183 cells containing the E1/E3-deleted pAdEasy Ad5 vector backbone **(Figure 1)** [32]. The E1 deletion renders subsequent adenoviruses replication deficient except in E1 helper cell lines (e.g. HEK293 cells). Transformed BJ5183 cells were incubated in SOC outgrowth medium (NEB, B9020S) for 1 hour at 200 rpm, and then plated onto Luria-Bertani (LB) agar containing 50 μg/ml kanamycin and 100 μg/ml ampicillin and allowed to grow at 37°C for 48 hours.

**Figure 1.**
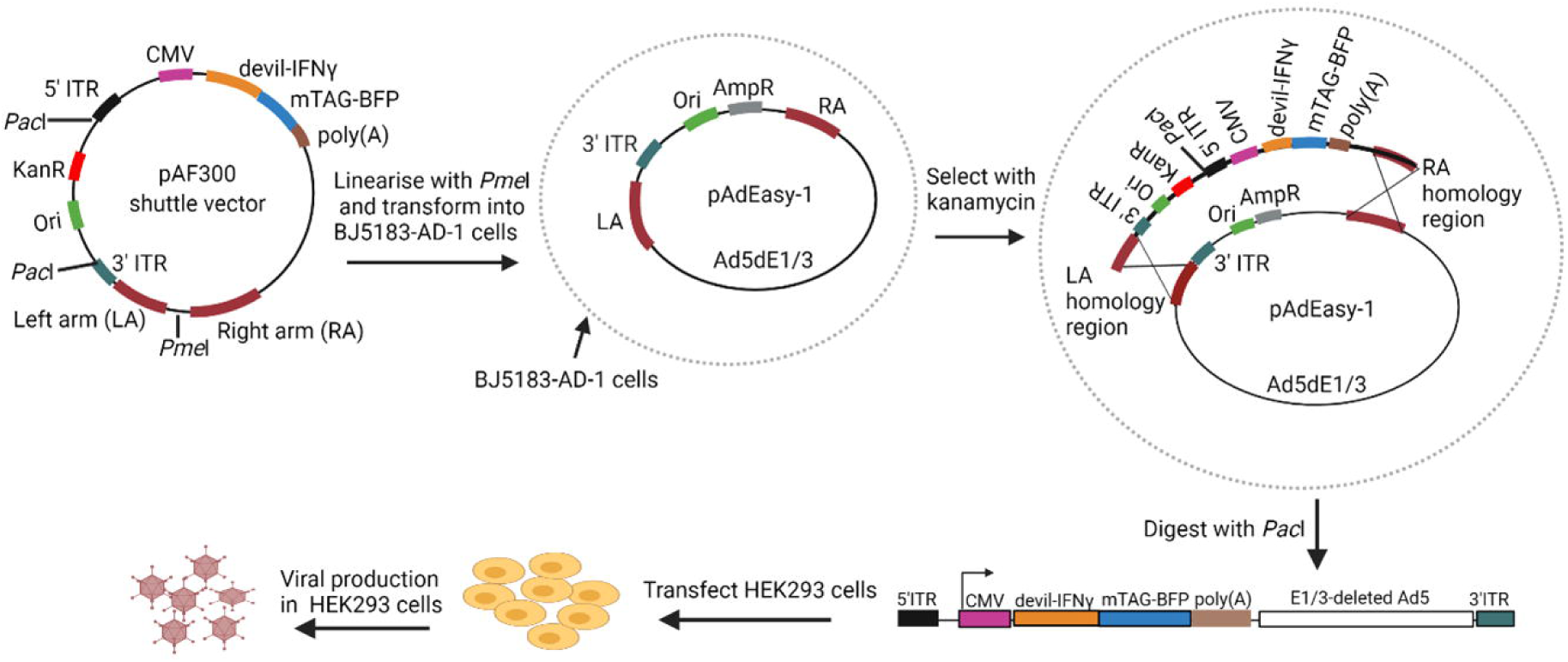
Schematic of the generation of the AdvAF300 adenoviral vector. The codon optimised devil IFN-γ gene fused to mTAG-BFP reporter was first cloned into a shuttle vector. The resultant plasmids were linearised by digesting with restriction endonuclease *Pme*I, and then transformed into BJ5183 bacterial cells pre-transformed with the Ad5 backbone vector, pAdEasy-1. Recombinant plasmids were selected for kanamycin resistance. Finally, the recombinant plasmids were linearized with *Pac*I and transfected into HEK293 cells for adenoviral vector packaging and production. Ad5dE1/3: Ad5 backbone without the E1 and E3 regions; CMV: cytomegalovirus promotor; ITR: inverted terminal repeat; LA: left arm; RA: right arm; poly(A): polyadenylation site. Figure created using Biorender.

BJ5183 cells with candidate plasmids were inoculated into 5 mL of LB broth with 50 μg/ml kanamycin and incubated overnight at 37°C and 200 rpm. A starter culture of 0.25 mL was then transferred to 250 mL of Luria broth and incubated overnight at 200 rpm. The plasmid DNA was extracted from the bacterial cultures using NucleoBond™ Xtra Midi Plus (Macherey-Nagel, 740412.10) according to the manufacturer’s guidelines. Recombinant plasmids (pAdvAF300) were then sent to the Core Sequencing Facility at the University of Technology Sydney for DNA sequencing.

### Production and testing of Ad5-IFN-γ

We performed an initial test of the replication-deficient Ad5-IFN-γ (see Supplementary Materials) prior to production of the master virus seed stock by the Vector and Genome Engineering Facility at the Children’s Medical Research Institute (Westmead, New South Wales, Australia). Briefly, the pAdvAF300 plasmid was linearised with *Pac*I (NEB, R0547S) and transfected into E1-complement HEK293A cells (ThermoFisher, R705-07) for the adenovirus packaging and generation. The Ad5 working virus seed stock was used to infect 20 × 150 mm plates for amplification of the virus. The cell lysate was harvested and the virus was purified using a Caesium chloride gradient and quantified using PCR as previously described [33]. The resulting master virus seed had a concentration of 1.93 × 10^12^ viral particles/mL and was stored at −80°C in 10% glycerol in phosphate-buffered saline (PBS).

### In vitro transduction of cell lines

The purified virus was tested *in vitro* in multiple cell lines at varying cell seeding densities, multiplicities of infection (MOI), and culture times in three different experiments. First, DFT2-SN, human A549, dog A-72, and Chinese hamster ovary (CHO) cells were seeded at the density of 1×10^5^ cells/well and incubated at 37°C and 5% CO_2_ in a humified incubator for 2 days before culture with experimental treatments for 96 hours. In the second experiment, DFT2-JV and DFT2-SN were seeded at 5 × 10^4^ cells/well and incubated for 5 days before culture with experimental treatments for 36 or 72 hours. In the third experiment, DFT1-1426, DFT1-4906, and devil FBB were plated at 5 × 10^4^ cells per well and cultured for 3 days before culture with experimental treatments for 36 or 72 hours.

In each case, the cells were seeded in 200 μL per well in flat bottom 96-well plates (Corning, 3599) in RPMI 1640 medium supplemented as above. Before treatment with the virus, the cells were washed with PBS (pH 7.4). Cells were treated with Ad5-IFN-γ at MOI of 1, 10, 100 or 1000 in a serum-free RPMI 1640 medium. Purified recombinant IFN-γ (pAF29) was used as a positive control at 10 ng/mL [34]. After 2 hours of incubation at 37°C and 5% CO_2_ the serum-free media was replaced with RPMI 1640 medium supplemented as above.

### Flow cytometry

Cells were rinsed with PBS (pH 7.4), enzymatically detached from the plate using TrypLE Express (Gibco, 1260-013), and transferred to V-bottom 96-well plates (Corning, 3894). Cells were then centrifuged at 500 × g for 3 min at 4°C and supernatant discarded. Cells were then stained with Live Dead near-IR fixable dye (Invitrogen, L34976) for determination of live cells. Cells were then blocked with 1% normal goat serum (Thermo Fisher Scientific, 01-6201) in flow cytometry buffer (PBS with 0.5% BSA, 0.05% NaN3) for 30 min at 4°C, followed by centrifugation at 500 × g for 3 min at 4^°^C and discarding supernatant. Cells were then incubated with 0.4 µL/sample of anti-devil β2m mouse antibody [6] for 30 min at 4°C. The cells were washed with flow cytometry buffer. Then 0.4 µg/sample of an Alexa Fluor 488-conguated goat anti-mouse IgG antibody (Invitrogen, A-11029) was added to the cells and incubated for 30 min at 4°C. Cells were then washed as above and then resuspended in FACS fixation buffer (PBS with 0.05% NaN3, 2% glucose and 0.4% formalin). Fluorescence intensity for β2m and BFP were detected at the wavelength of 488/530 and 405/450 nm respectively, using BD FACSCanto II flow cytometer (BD Biosciences, San Jose, CA, United States). All flow cytometric analysis was conducted using FCS Express (6th Edition).

## Results

To initially test for functional effects of IFN-γ expressed in Ad5-IFN-γ treated cells, we incubated DFT1-C5065 with either 10 µg/mL of purified IFN-γ or crude supernatant from Ad5-IFN-γ transfected HEK293A cells. DFT1 cells were gated for singlets and sub-gated by forward and side scatter. Incubation of DFT1 cells with purified devil IFN-γ or Ad5-IFN-γ supernatant resulted in similar upregulation of β2m (**Figure S1**). Following this functional confirmation of the Ad5-IFN-γ adenoviral construct we produced a purified master virus seed stock of Ad5-IFN-γ. We used transmission electron microscopy (TEM) to visually assess the morphology of the Ad5-IFN-γ master virus seed. TEM Imaging at 200,000x showed the characteristic icosahedral structure and size of the adenoviruses (**Figure S2**).

To evaluate the ability for a human adenovirus to transduce devil cell lines and express functional transgenes, we treated DFT2-SN, A549, A-72, and CHO-K1 cells with the Ad5-IFN-γ master virus seed. IFN-γ and Ad5-IFN-γ treatments increased cell death, so we gated on live cells only for our analyses (**Figure 2A**). β2m was strongly upregulated on DFT2-SN cells in wells treated with the purified recombinant devil IFN-γ. Expression of β2m on DFT2-SN cells increased with increasing MOI (**Figure 2B**).

**Figure 2.**
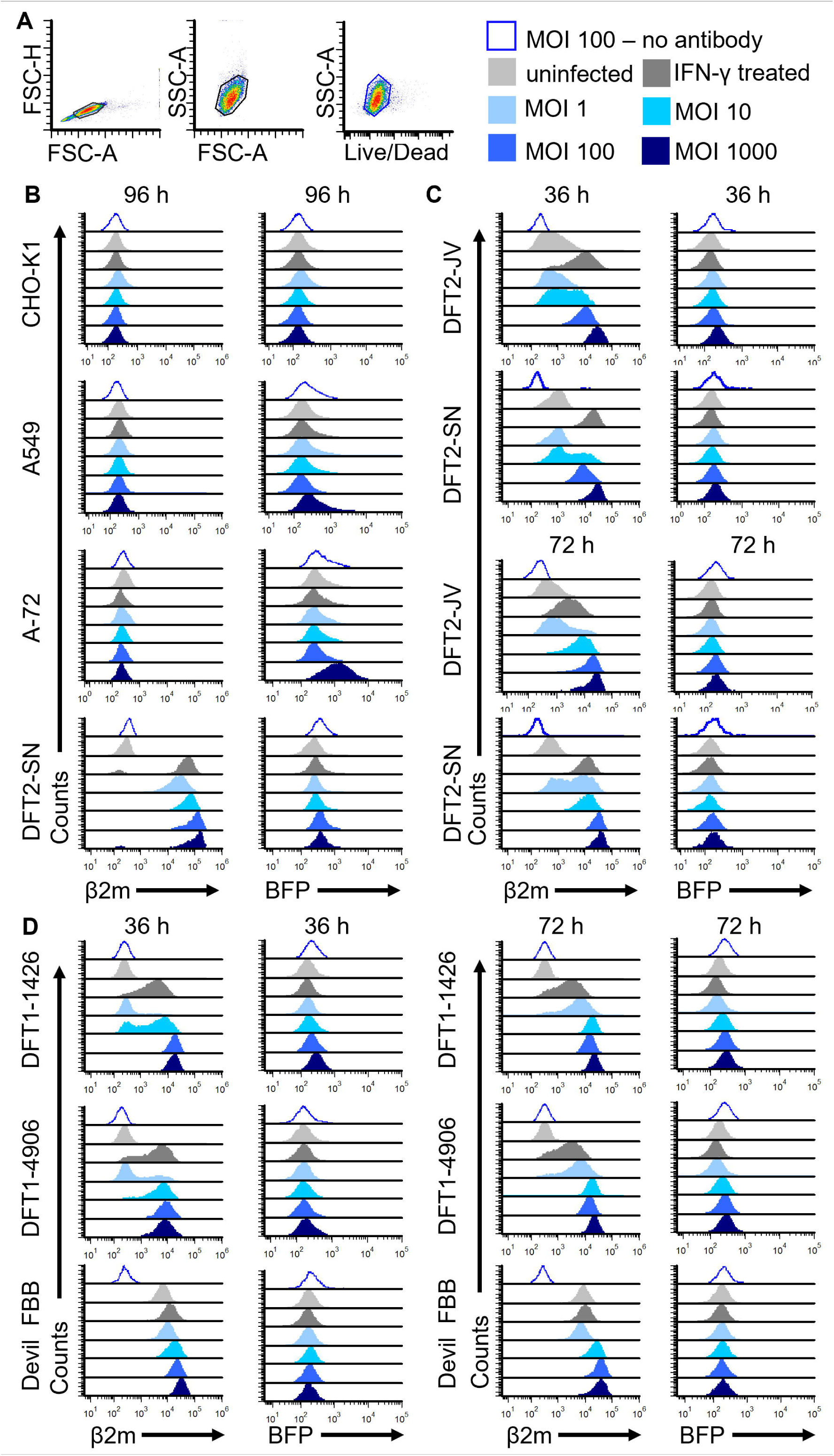
Adenoviral-expressed IFN-γ upregulates β2m on devil cells. (A) Vating strategy for single cells, forward scatter x side scatter, and live/dead. (B) β2m and BFP fluorescence intensities in CHO-K1, A549, A-72 and DFT2-SN cells harvested 96 hours after treatment with IFN-γ or Ad5-IFN-γ or no treatment. (C) DFT2-JV and DFT2-SN cells harvested at 36- and 72-hours post-treatment. (D) DFT1-1426, DFT1-4906, and devil fibroblasts (FBB) cells harvested at 36- and 72-hours post-treatment. Surface β2m was assessed using mouse anti-devil β2m antibody followed by goat anti-mouse IgG conjugated to Alexa Fluor 488 before they were analyzed by flow cytometry. For B-D the Y-axis represents cell count, X-axis represents biexponential scale of fluorescence at 488/530 nm (ex/em) for β2m and at 405/450 nm (ex/em) for BFP. The legend shows cells treated with MOI 100 but not stained with antibodies in open histograms, untreated cells with antibody in light gray, recombinant IFN-γ treated cells in dark gray, and Ad5-IFN-γ treated cells with MOI ranging from 1 to 1000 are shown in light blue to dark blue. Plots are representative of n=3 technical replicates for each treatment.

A549 cells are known to be permissive to transduction by Ad5, whereas CHO cells are reported to be non-permissive. As expected, the devil-specific anti-β2m monoclonal antibody did not bind to CHO-K1, A-72, and A549 cells (**Figure 2B**). However, BFP fluorescence increased in A549 and A-72 cells but not in CHO cells, suggesting Ad5-IFN-γ entered A549 and A-72 and expressed CMV-driven transgenes. BFP-expression generally increased with increasing MOI, but was absent in CHO cells, untreated cells, and cells treated with recombinant IFN-γ. There appeared to be a marginal increase of BFP in DFT2-SN cells at higher MOI.

The next experiment used DFT2-SN and DFT2-JV cell lines at two earlier time points than experiment one. Recombinant IFN-γ treatment again resulted in strong upregulation of β2m on DFT2 cells (**Figure 2C**); upregulation of β2m on Ad5-IFN-γ treated cells again increased with increasing MOI. BFP expression increased slightly at higher MOI.

We next tested Ad5-IFN-γ on DFT1-1426, DFT1-4906, and devil fibroblasts (FBB). β2m expression on DFT1 cells strongly increased with increasing MOI (**Figure 2D**). FBB constitutively express high levels of β2m, but expression increased with increasing MOI. BFP expression was again low in all cell lines but appeared to increase slightly at higher MOI in DFT1 cells.

## Discussion

Human Ad5 has been widely used as a gene delivery and vaccine platform for humans and other eutherian mammals. However, it was unknown if Ad5 could transduce marsupial cell lines and express transgenes. Here we report that a replication-deficient Ad5 vector has ability to transduce both normal and tumour cells from the Tasmanian devils and express functional transgenes using a CMV promotor. The ability to immunotherapies and vaccines *in vivo* in an endangered species is extremely limited, so this *in vitro* study is an important milestone towards improved immunotherapies and vaccines for devil conservation. Additionally, this opens the door to adenoviral-based vaccines for other marsupial diseases.

Our results using IFN-γ expressed via a human adenovirus agree with previous studies showing that IFN-γ strongly upregulates β2m on devil facial tumour cells [6, 35, 36]. More recently, genetically modified DFT1 cells that either constitutively upregulate β2m or have β2m knocked out have been used to conclusively show that MHC-I is a major target in immunotherapy-induced DFT1 regressions and rare cases of natural DFT1 regressions [7, 10, 37, 38]. Our results here using the Ad5-IFN-γ vector open the door to relatively low-cost immunotherapy to potentially treat devils with DFT1 or DFT2.

The functional effects of IFN-γ were confirmed via upregulation of β2m on transduced and bystander devil cells; the lack of β2m on human, dog, and hamster cells was expected due to using a devil specific anti-β2m monoclonal antibody and the low probability of devil IFN-γ binding to IFN-γ receptors in other species. However, the relatively low expression of BFP adenovirus-permissive cell lines, including human A549 and dog A-72 cells, was surprising. One possibility is that the IFN-γ-BFP fusion is rapidly secreted and thus does not accumulate in transduced cells to yield a strong fluorescent signal, whereas IFN-γ-induced upregulation of β2m can be detected on the cell surface for several days.

The low fluorescence intensity of BFP could also be due to rapid photobleaching of BFP compared to other traditional fluorescence reporters such as the green fluorescence protein [39– 41]. Perhaps most interestingly, the BFP in the Ad5-IFN-γ contains an Leu-Val-Gly-Gly-Ser amino acid sequence that is a cleavage site for the cysteine protease encoded in the Ad5 genome that is necessary for adenoviral maturation [42, 43]. Previous work has shown the fluorescent proteins fused to adenoviral structural proteins are degraded by the adenoviral protease [44]. It is possible that BFP may have been degraded by the protease, resulting in low BFP signal.

An effective immunotherapy would be a helpful tool for devils with DFTD, but a more valuable conservation tool would be a prophylactic vaccine that can prevent DFTD. IFN-γ stimulated whole DFT1/2 cell vaccines provide the full spectrum of potential tumour antigens, whereas prophylactical viral vector-based vaccine would need to encode tumour antigens into the vector. For human cancers this has historically been a major challenge because each cancer is different. The problem of identifying effective tumour-specific antigens or tumour-associated antigens is more tractable because DFT1 and DFT2 are transmissible cancers that carry ancestral genome mutations forward to new hosts. Thus, tumour antigens for use in prophylactic vaccines need to be identified for only two different tumours, rather than new antigens for each new host.

## Conclusion

Human Ad5 can transduce and express functional transgenes in devil cell lines. This warrants further development of human adenoviral vectors for use as prophylactic vaccines and immunotherapies for DFTD and potentially other marsupial diseases. Future studies will be needed to assess the *in vivo* potential of the adenoviral vector to induce an anti-viral immune response and stimulate antigen processing and presentation in healthy and diseased devils.

## Authors and contributors

ABL, ALP, ANK, ASF, GSL, and LL conceived and designed the study. ASF, ANK, and JMD performed the experiments. ABL, ALP, ANK, and ASF analysed the data. ANK and ASF wrote the manuscript, and all others edited the manuscript.

## Data availability

Data related to this paper will be publicly available through the University of Tasmania Research Data Portal.

## Ethical statement

The research study was approved by the University of Tasmania’s Institutional Biosafety Committee (UTAS-38-2016).

## Acknowledgments

We thank Dr Hannah Siddle for providing us with the anti-β2m monoclonal antibodies and Dr Terry Pinfold for help with flow cytometry. We wish to thank Dr Olivier Bibari for his assistant with transmission electron microscopy and processing of the viral vector image. We thank Prof Greg Woods and the devil immunology group for help with the research. We thank Predrag Kalajdzic for help preparing the virus stock.

