## Supplementary Materials for "A human adenovirus encoding IFN-γ can transduce Tasmanian devil facial tumour cells and upregulate MHC-I"

**Title**

*Initial testing of virus supernatant*

Five µg of the pAdvAF300 plasmid was linearised with *Pac*I (NEB, R0547S). To produce the replication-deficient Ad5-IFN-γ, the linearized plasmids were electroporated into E1-complement HEK293A (Thermo Fisher Scientific, R705-07) using a SF Cell Line 4D-Nucleofector™ X Kit L (Lonza, V4XC-2012) and pulse CM-130 setting as previously described [30]. The electroporated HEK293 cells where then transferred to a T25 flask and cultured in RPMI 1640 medium. Following confirmation of BFP expression and cytopathic effects, the cells were lysed using two freeze/thaw cycles in a dry ice and ethanol bath and used to transduce HEK293A cells in a T75 flask to produce a larger crude Ad5-IFN-γ lysate. To test if functional IFN-γ was secreted from the infected cells, the supernatant was collected and added to DFT1-C5065 cells. Purified recombinant IFN-γ (pAF29) was used as a positive control at 10 ng/mL. The cells were incubated overnight and BFP and surface β2m expression were analysed via flow cytometry.

*Transmission Electron Microscopy*

Transmission electron microscopy (TEM) was used to visualise the icosahedron structure of the adenovirus in the purified virus stock. Briefly, TEM grids were coated with poly-L-lysine for 15 min before adding the virus sample by placing the grid face down on a drop (~2.5 uL) of the virus master seed stock. The virus absorbed for 1 minute. Extra solution was removed by using filter paper and the grid left to dry in air at room temperature. Grids were negatively stained with 1% Uranyl acetate solution in 50% ethanol. The grids were then rinsed in water for 1 minute and spot-dried with filter paper. Finally, grids were allowed to air dry and then imaged using a TEM FEI Tecnai G2 electron microscope.

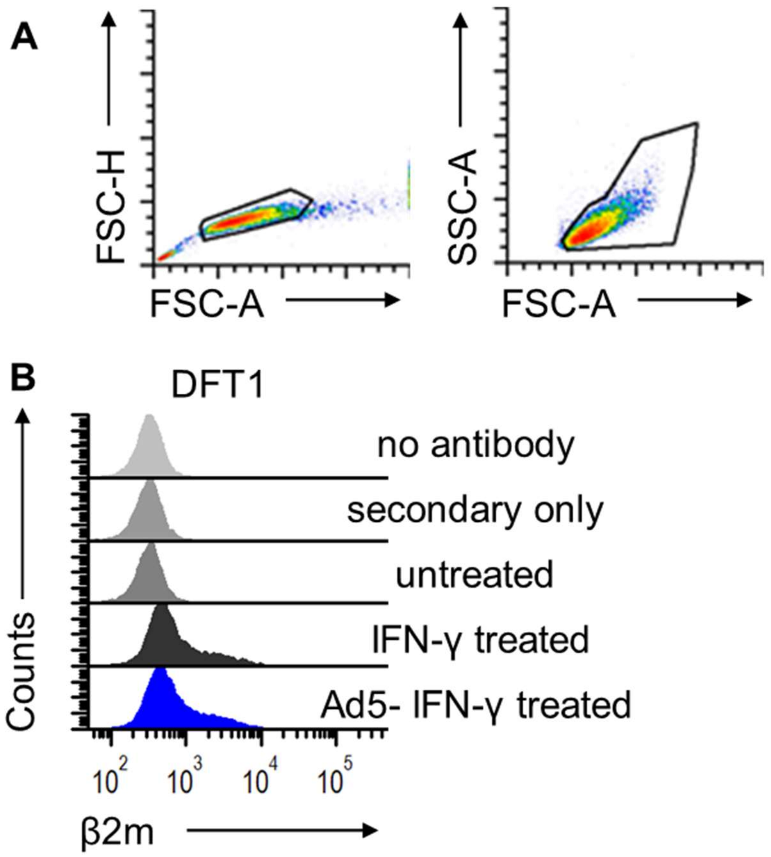


**Figure S1.** Upregulation of β2m on the DFT1 cells using supernatant from Ad5-IFN-γ transduced HEK293. (A) Gating strategy and (B) flow cytometric analysis of β2m expression on DFT1-C5065 cells treated with recombinant IFN-γ or Ad5-IFN-γ supernatant. Cells treated with 10 µg/mL recombinant IFN-γ were used as positive controls. Cells were stained with mouse anti-devil β2m antibody followed by goat anti-mouse IgG conjugated to Alexa Fluor 488 and then analyzed by flow cytometry.

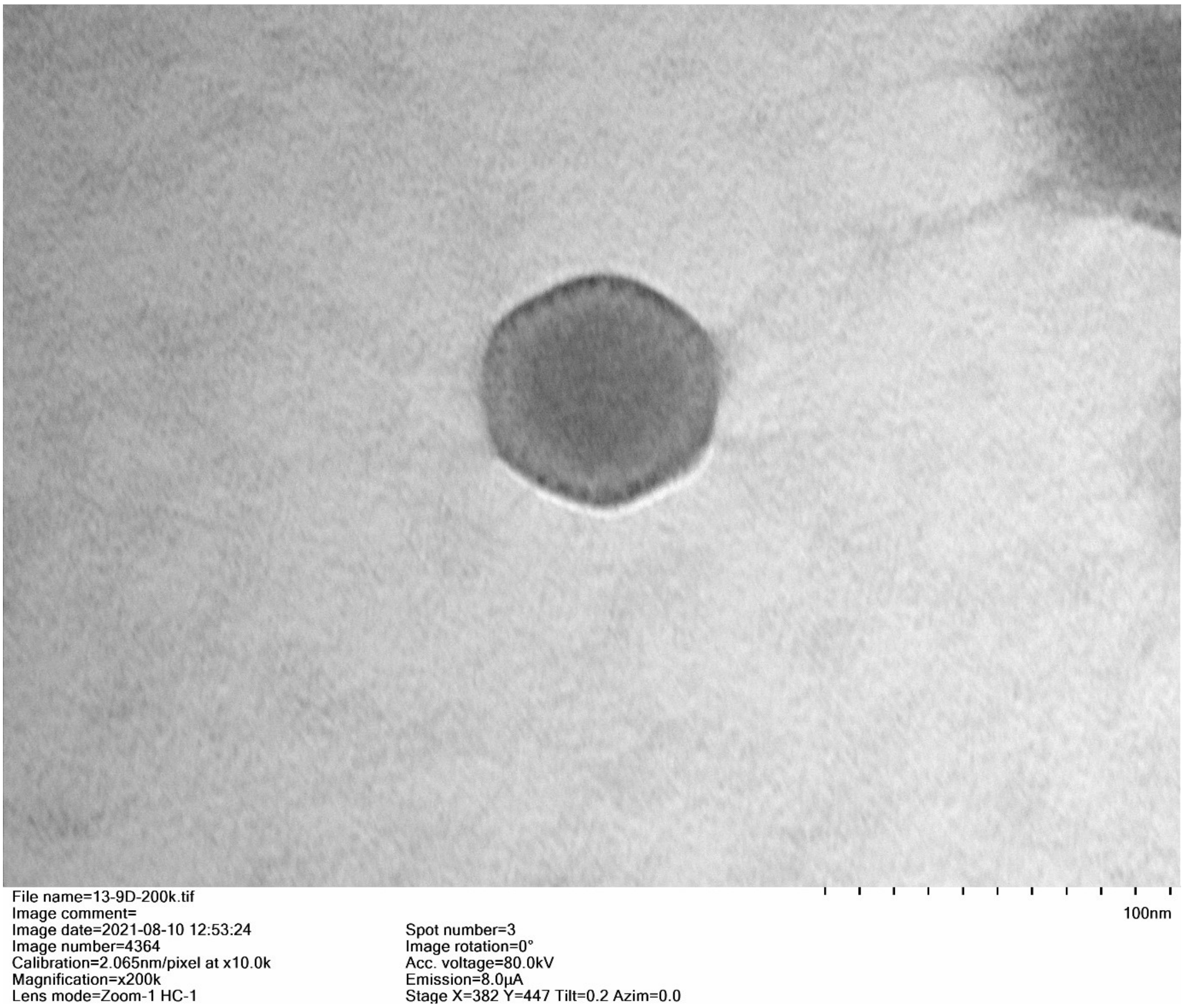


**Figure S2**. Transmission electron microscopy image of the Ad5-IFN-γ. The morphology is characteristic of icosahedral adenoviruses. Scale shows 100 nm; magnification of the image: 200,000x.
